# Integrative multispecies omics reveals a hierarchy of cold-responsive regulatory network launched by circadian components in rosids

**DOI:** 10.1101/2022.10.03.510673

**Authors:** Liangyu Guo, Zhiming Xu, Shuo Wang, Yuqi Nie, Xiaoxue Ye, Xuejiao Jin, Jianhua Zhu, Wenwu Wu

**Affiliations:** State Key Laboratory of Subtropical Silviculture, School of Forestry and Biotechnology, Zhejiang A&F University, Lin’an, Hangzhou 311300, China; Institute of Tropical Biosciences and Biotechnology, Chinese Academy of Tropical Agricultural Sciences, Haikou 571101, Hainan, China; School of Life Sciences, Anhui Agricultural University, Hefei, 230036, China; Department of Plant Science and Landscape Architecture, University of Maryland, College Park, Maryland 20742, USA

**Author notes:** Correspondence should be addressed to X.J, J.Z. or W.W. (, Dr. Wu is fully responsible for the distribution of the materials associated with this article). Equal contributors (Liangyu Guo, Zhiming Xu, Shuo Wang).

**Keywords:** transcription factor, RNA binding protein, circadian clock, hierarchical network, cold stress, rosids

## Abstract

Elucidating regulators and molecular mechanisms underlying gene transcriptional and post-transcriptional co-regulatory network is key to understand plant cold-stress responses. Previous studies were mainly conducted on single species and whether the regulators and mechanisms are conserved across different species remains elusive. Here, we selected three species that diverged at early evolution of rosids (93–115 million years ago) and integrated phylotranscriptome and ChIP/DAP-seq datasets to identify cold-responsive regulators and their regulatory networks. First, we found over ten thousand cold-responsive genes including differentially expressed genes (DEGs) and alternative splicing genes (DASGs) in each species. Among the DEGs, genes encoding a set of transcription factors (TFs) (AP2/ERF, MYB, WRKY, NAC, etc.) and RNA binding proteins (RBPs) (Ribosomal, RRM, DEAD, Helicase_C, etc.) are conserved in cold responses in rosids. Compared to TFs, RBPs show a delayed cold-responsive pattern, likely suggesting a hierarchical regulation of DEGs and DASGs. Between DEGs and DASGs, we identified 259 overlapping DE-DASG orthogroups and interestingly, pathway analysis on each dataset of DEGs, DASGs, and DE-DASGs coincidently shows an enrichment of circadian rhythm. Evidentially, many circadian components are cold-regulated at both transcriptional and post-transcriptional levels. Moreover, we reasoned 226 cold-responsive genes regulated by at least two of five circadian components (CCA1, LHY, RV4, RVE8, and RVE7) in rosids. Finally, we unveiled a conserved hierarchical network in dynamic transcriptional and post-transcriptional regulation of cold-responsive genes launched by circadian components in rosids. Together, our results provide insights into core regulators and mechanisms underlying cold-responsive regulatory network across rosids, despite a long evolutionary history.

## Introduction

For adaptation to adverse environmental conditions, angiosperms have evolved sophisticated and advanced mechanisms and become the largest land plant lineage with more than 350,000 known species widely distributed from tropical to polar terrestrial zones^1^. Among the adverse environmental conditions, seasonal and daily variation of temperature is a fundamental component that affects plant growth, productivity, and geographical distribution^2,3^. Under cold stress, plants undergo physiological and molecular changes that increase the levels of specific proteins, metabolites, and phytohormones, which are generally regulated by transcription factors (TFs) and RNA-binding proteins (RBPs) at transcriptional and post-transcriptional levels^3-6^.

Thousands of differentially expressed genes (DEGs) were identified involved in cold stress^7-9^ and many TFs have been suggested in regulating and managing their expression^3,4,10^. The best characterized TFs are C-repeat binding factor (*CBF*) genes that are rapidly and transiently induced by cold stress and can enhance freezing tolerance of plants through binding to *cis*-elements (CCGAC) in the promoters of cold-responsive (*COR*) genes^11,12^. RNA-seq analysis of *Arabidopsis cbf123* triple mutant showed that hundreds of *CORs* are regulated through a CBF-dependent pathway^13,14^. Activation of CBF target *CORs*, such as *COR15A, COR47, KIN1*, and *RD29A*, leads to an increase in plant freezing tolerance^12,15,16^. In addition, cold responsiveness of plants is regulated by circadian clock system, which is involved in plant stress responses through controlling daily energy expenditure and enabling plants to grow and reproduce under seasonal and daily temperature variation^17-21^. In the clock system, some circadian clock-related TFs, such as CIRCADIAN CLOCK ASSOCIATED 1 (CCA1), LATE ELONGATED HYPOCOTYL (LHY), and REVEILLES (RVEs), have been proven involved in cold tolerance by directly regulating the expression of downstream *CORs*, such as *CBFs, COR15A*/*15B, COR28*, and *COR27*^2,22-24^.

On the other hand, post-transcriptional regulation is also a key process in reprogramming gene expression for plant to tolerate or survive constantly changing in temperature^3,4^. Similar to TFs in regulating gene transcription, RBPs are essential in controlling post-transcriptional RNA processing, including capping, polyadenylation, alternative splicing (AS), RNA export, translation, and turnover^7,25^. Under cold stress, upregulation and/or downregulation of RBP genes may cause differentially post-transcriptional RNA processing. During RNA processing, AS is an important post-transcriptional mechanism by producing different transcripts and possibly protein variants from a single gene for increasing transcriptome and proteome diversity^26^. The concentration, localization, and activity of RBPs determine AS events to produce different transcripts^27,28^. In plants, extensive AS events have been implicated in various abiotic stresses, including cold, salt, heat and other environmental stresses^7,8,29,30^. For example, genome-wide AS profiling in *Arabidopsis* showed a massive differentially alternative splicing genes (DASGs) after cold treatment and suggested that plants may fine-tune AS events of gene transcripts for adapting to cold stress^7^. Among DASGs, *CCA1*, a core component of circadian clock, has two splice variants: CCA1α and CCA1β. CCA1β competitively inhibits the DNA binding activity of CCA1α and LHY by forming nonfunctional heterodimers of CCA1β-CCA1α and CCA1β-LHY^31^. Under cold stress, CCA1β production is repressed to release the activity of CCA1α and LHY, whereby CCA1α and LHY promote the expression of downstream *CORs* for improving plant cold tolerance^31^.

In the past years, numerous cold-induced DEGs and DASGs were identified in different plant species by high-throughput methods^7,8,29,30,32^. Identifying core regulators (e.g., TFs and RBPs) and elucidating molecular mechanisms underlying the outcome of DEGs and DASGs are key to understand the reprogramming of gene regulatory network in plant response to cold stress. The previous studies were generally performed on single plant species, especially the reference model *Arabidopsis thaliana*, and whether the regulators and mechanisms underlying transcriptional and post-transcriptional changes are conserved in different plant lineages remains elusive. In this study, we focused on rosids, a large group of eudicots in angiosperms, and selected three plant species: *Carya illinoinensis* and *Populus trichocarpa* from fabids (rosid I) and *Arabidopsis thaliana* from malvids (rosid II). Fabids and malvids are two main clades of rosids, comprising of >40,000 and ∼15,000 species, respectively^33^. The three plant species diverged during early evolution of rosids and evolved independently for over 100 million years^34^. The long evolutionary history caused great differences in their growth habits and geographic distributions, and accordingly, we asked whether the regulators and mechanisms underlying gene transcriptional and post-transcriptional regulation are conserved across rosids in response to cold stress.

To solve the question, we performed omics analysis of genomes, transcriptomes, and regulomes and identified common core regulators in regulating cold responses across the three plant species. First, by RNA-seq experiments, we investigated global expression profiles of cold-induced DEGs and DASGs in each of the three studied species. Subsequently, we analyzed cold responses of TFs and RBPs and identified those TF and RBP families that are conserved in cold responses, which likely contribute to DEGs and DASGs in rosids. Moreover, we explored overlapping genes (i.e., DE-DASGs) of DEGs and DASGs and performed homologous analysis to identify conserved DE-DASG orthogroups in rosids. Finally, we selected five circadian components and integrated transcriptomes and regulome maps to identify their target *CORs* for constructing a hierarchical cold-responsive regulatory network.

## Results

### Global gene expression profiles in rosids under cold stress

Physiological and molecular changes in plants under cold stress are usually caused by up- or downregulated expression of genes^3^. To capture the landscape of DEGs under cold stress, we performed RNA-seq experiments in three rosid species before and after cold treatments (**see Methods and materials**). Based on RNA-seq analysis, we identified a huge number of cold-induced DEGs: 7,579, 8,878, and 8,480 respectively in *A. thaliana, C. illinoinensis*, and *P. trichocarpa* (**Supplementary Table 1**). Notably, more upregulated than downregulated DEGs are observed in all three plants (**Fig 1a**). Expression profiles show that these upregulated DEGs are mainly clustered in three expression patterns following three timepoints of cold treatments: 2 hours (h), 24 h, 168 h after cold treatments (**Fig 1b**). Inspecting the expression patterns, we found that the number of upregulated DEGs in *A. thaliana* is significantly more at 2 h but less at 168 h of cold stress treatment than the counterparts in other two species. Likewise, there are significantly more downregulated DEGs at 2 h and less downregulated DEGs at 168 h in *A. thaliana* than those in *C. illinoinensis* and *P. trichocarpa* (**Fig 1b**). These findings reveal that over seven thousand genes are differentially cold-regulated in rosids with three common expression patterns but likely in different cold-responsive speeds: an early extensive response in *A. thaliana* but a delayed extensive response in *C. illinoinensis* and *P. trichocarpa*.

**Fig 1.**
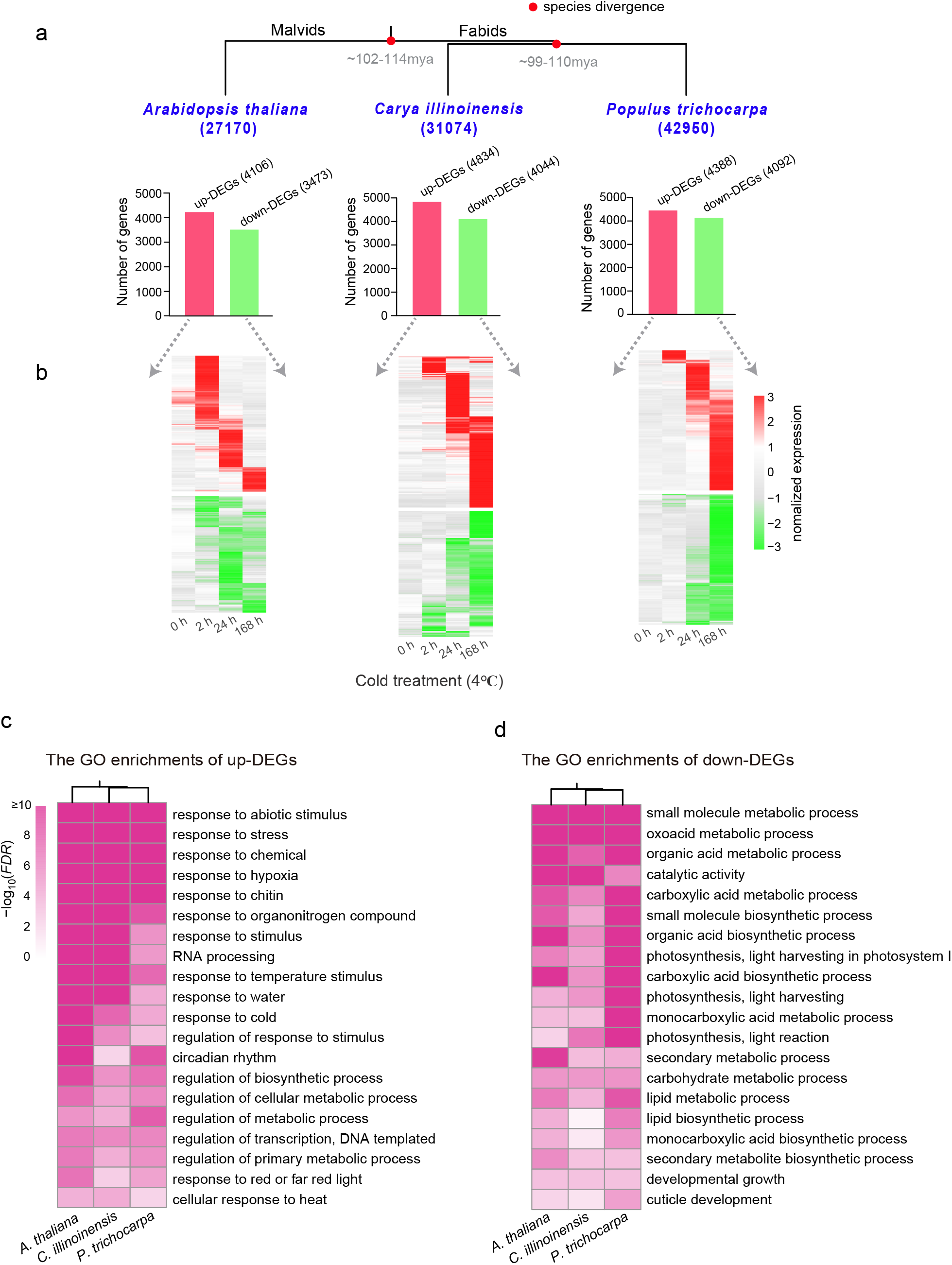
Identification and enrichment analysis of cold-induced differentially expressed genes (DEGs). (a) Overview of DEGs under cold stress. The number in parentheses below each species indicates the number of all coding genes in the genome. The evolutionary relationship and divergence times of the species were obtained from the website TimeTree (Kumar et al., 2017). Bar heights indicate gene numbers of upregulated DEGs (in red) and downregulated DEGs (in green). (b) Expression profiles of DEGs before and after cold treatments (4¼). The expression values of DEGs at four different timepoints (0, 2, 24, and 168 h) of cold treatments were normalized by row scale. (c-d) GO term enrichments of upregulated DEGs and downregulated DEGs. The top 20 significant GO terms were shown in a conserved manner with common cold-responsive biological processes in three species.

### Common biological processes of cold responses in rosids

Despite of different cold-responsive speeds between the three plant species, Gene Ontology (GO) term enrichment analysis of the DEGs showed a very similar enrichment of biological processes (**Fig 1c, 1d**). The upregulated DEGs are mainly enriched in abiotic stress processes, e.g., response to hypoxia, water, and cold, and circadian rhythm (**Fig 1c**). Additionally, regulations of transcription and RNA processing are also amongst the most enriched GO terms, which suggests that the regulators (e.g., TFs and RBPs) at transcriptional and post-transcriptional level are likely to be more activated under cold stress (**Fig 1c**). The downregulated DEGs are mainly enriched in processes such as catalytic activity, photosynthesis, developmental growth, small molecular and/or lipid biosynthetic/metabolic process (**Fig 1d**). It is reasonable that for improving cold tolerance, plants need to reduce expression of the genes involved in activities of plant development and growth^3^.

### Conserved cold-responsive TF families in rosids

Differential changes of gene expression at transcriptional level are likely caused by fluctuations of TFs^3,7^. To detect the roles of TFs under cold stress, we first identified a total of 1,717, 1,899, and 2,466 TFs in *A. thaliana, C. illinoinensis*, and *P. trichocarpa* genomes, respectively (**Supplementary Table 2, see Methods and materials**). Transcriptome analysis shows that over five hundred TFs are differentially cold-regulated, covering a high proportion of the genome TFs, that is, 32%, 35%, and 22% respectively in *A. thaliana, C. illinoinensis*, and *P. trichocarpa* (**Fig 2a, Supplementary Table 2**). Notably, there are more upregulated than downregulated TFs and their expression patterns in cold-responsive speeds agree well with the aforementioned global expression patterns of DEGs (**Fig 1b, Fig 2b**). This finding suggests more upregulated TFs might be involved in modulating differentially regulated genes under cold stress.

**Fig 2.**
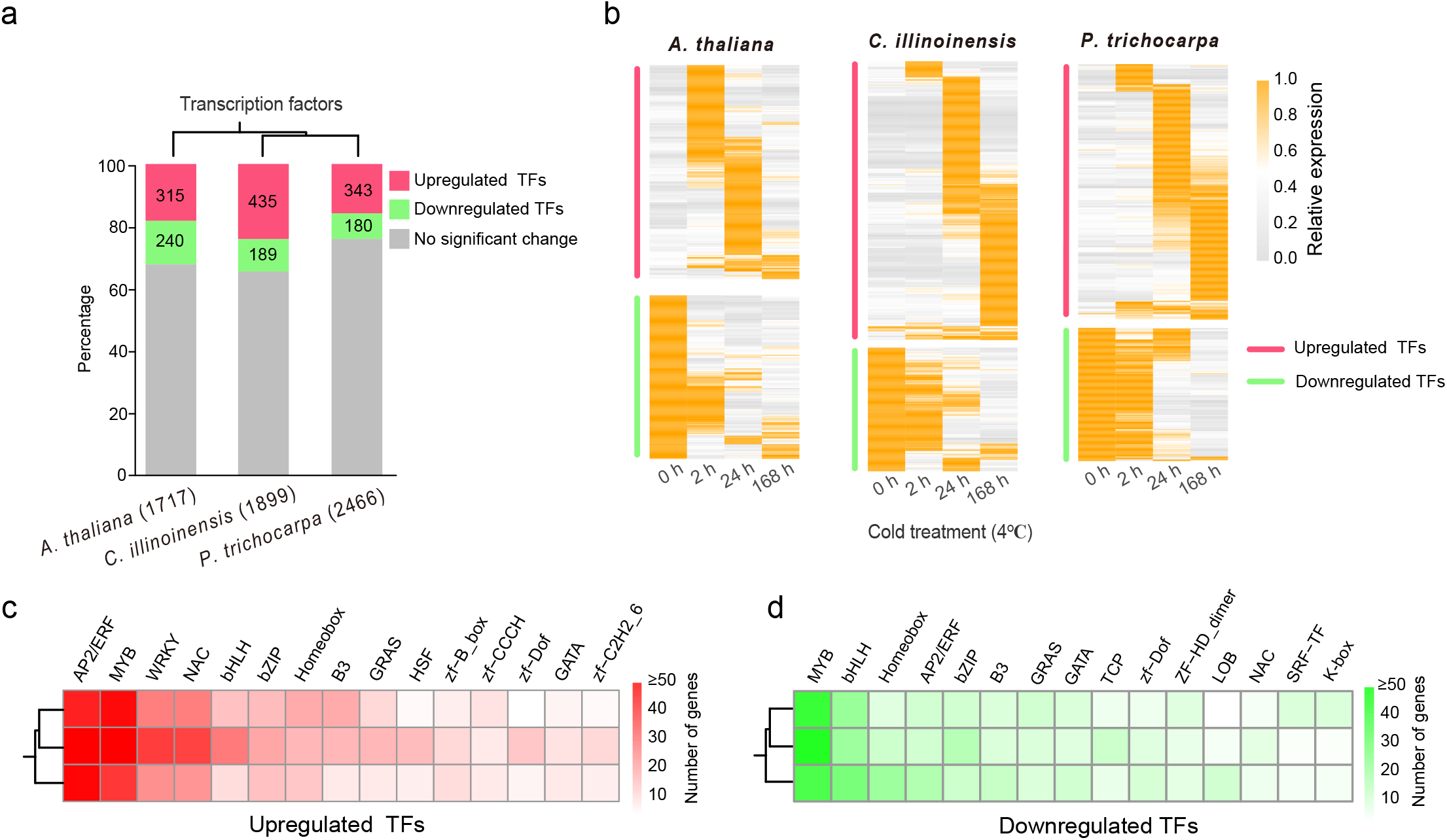
Cold-responsive TFs in conserved TF families. (a) Upregulated and downregulated TFs under cold stress. Bar percentage indicates the proportion of upregulated, downregulated, and no significant expression changed TFs to all TFs. The number in parentheses after the species indicates the number of TFs in the genome. (b) Expression profiles of up- and downregulated TFs under cold stress. The expression values of the TFs at four different timepoints (0, 2, 24, and 168 h) of cold treatments were normalized by row scale. (c-d) The numbers of up- and downregulated TFs in TF families. The top 15 up- and downregulated TF families were selected for visualization.

Moreover, we classified all TFs into gene families based on their protein domain architectures (**Supplementary Table 2, see Methods and materials**). Interestingly, the up- and downregulated TFs are respectively clustered in common gene families in three plant species (**Fig 2c, 2d)**. For example, the main families of upregulated TFs are AP2/ERF, MYB, WRKY, and NAC, which are the best-known TF families involved in abiotic stresses^35,36^. The counterparts of downregulated TFs are MYB, bHLH, Homeobox, and AP2/ERF. Notably, AP2/ERF and MYB are two most prominent TF families with many more upregulated members but also a considerable number of downregulated members in all three plants under cold stress. It is explainable that members from the same TF families can often function in a negative feedback mechanism in abiotic responses^37,38^. According to the maximum parsimony principle, we suggest that the conserved cold-responsiveness of those TF families in three plant species is largely descended from their common ancestor.

### Global alternative splicing profiles in rosids under cold stress

In addition to transcriptional regulation, we identified an enrichment of post-transcriptional regulation, such as RNA processing, in response to cold stress (**Fig 1c**). Based on transcriptome analysis, we detected 2,572, 7,428, and 7,817 cold-induced DASGs, which occupy 10%, 24%, and 18% of coding genes in *A. thaliana, C. illinoinensis*, and *P. trichocarpa*, respectively (**Fig 3a, Supplementary Table 3**). Among DASGs, many have multiple AS sites, producing 3,192, 11,972, and 13,373 differentially AS events (DASEs), respectively in *A. thaliana, C. illinoinensis*, and *P. trichocarpa* (**Fig 3a, Supplementary Table 3**). This is mainly caused by AS occurrences at the same or different exon/intron boundaries across a gene, which leads to more diversities of RNA transcripts. In consistence with previous studies^7,30^, intron retention (IR) is the most common AS mode in three plant species (**Fig 3b**). Despite a relatively small role of skipping exon (SE) in *A. thaliana*, SE provides a crucial role in other two plants (*C. illinoinensis* and *P. trichocarpa*). Further, GO term enrichment analysis shows that these DASGs from three different plant species are enriched in very similar biological processes, such as metabolic processes, gene expression, RNA splicing, and circadian rhythm (**Fig 3c**). Given a key role of RBPs in RNA processing^7,25^, thousands of cold-induced DASEs might at least partially result from fluctuations of RBP genes.

**Fig 3.**
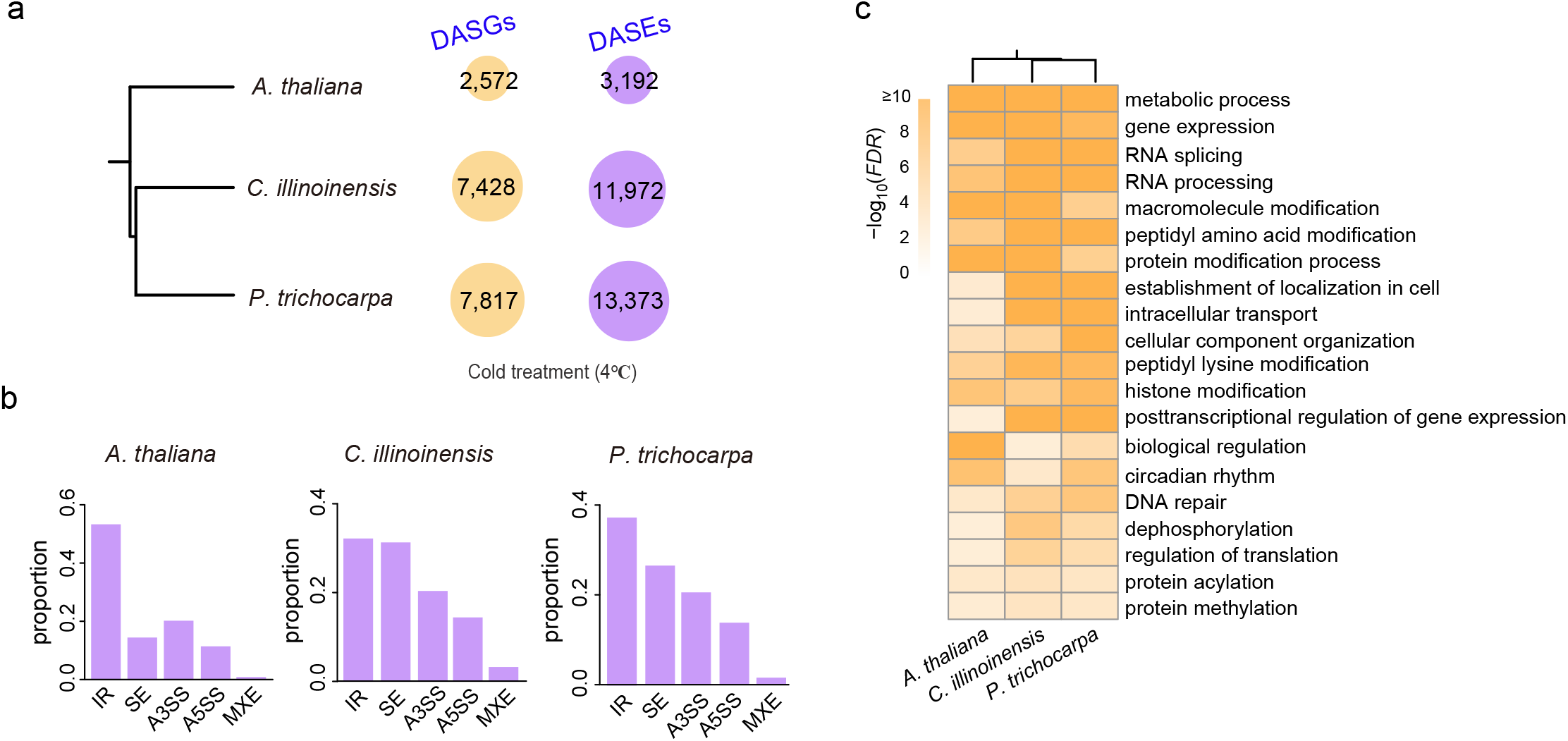
Identification and enrichment analysis of cold-induced alternative splicing genes (DASGs). (a) Overview of DASGs and DASEs under cold stress. The number in circle indicates the number of cold-induced DASGs or DASEs in the corresponding species. (b) Proportions of five AS types of cold-induced DASEs. IR: intron retention; SE: skipping exon; A3SS: alternative 3’ splicing site; A5SS: alternative 5’ splicing site; MXE: mutually exclusive exons. (c) GO term enrichments of DASGs. The top 20 significant GO terms were shown in a conserved manner with common cold-responsive biological processes in three species.

### A delayed cold responsivity of RBP genes

In *A. thaliana*, we collected a total of 788 high-confident RBPs that were experimentally confirmed by RNA-binding assays^25,39^. Using the RBPs for homologs searching, we identified 1,110 and 1,379 RBPs in *C. illinoinensis* and *P. trichocarpa*, respectively (**Fig 4a, Supplementary Table 4**). Transcriptome analysis shows that 36%, 31%, and 35% RBPs are differentially regulated by cold stress in *A. thaliana, C. illinoinensis*, and *P. trichocarpa*, respectively (**Supplementary Table 4**). Like TFs, we identified more upregulated RBPs than downregulated ones, especially in *A. thaliana* (**Fig 4a**). Interestingly, in contrast to TF genes, RBP genes show a delayed responsive pattern after cold treatment in all three plant species (**Fig 2b, 4b**). For example, only a relatively small number of RBP genes are observed to be differentially regulated at early stage (2 h) of cold stress. This finding shows that cold responsivity of RBP genes lags after that of TF genes, which might suggest a hierarchy of temporal dynamics of transcriptional and post-transcriptional regulatory network in response to cold stress.

**Fig 4.**
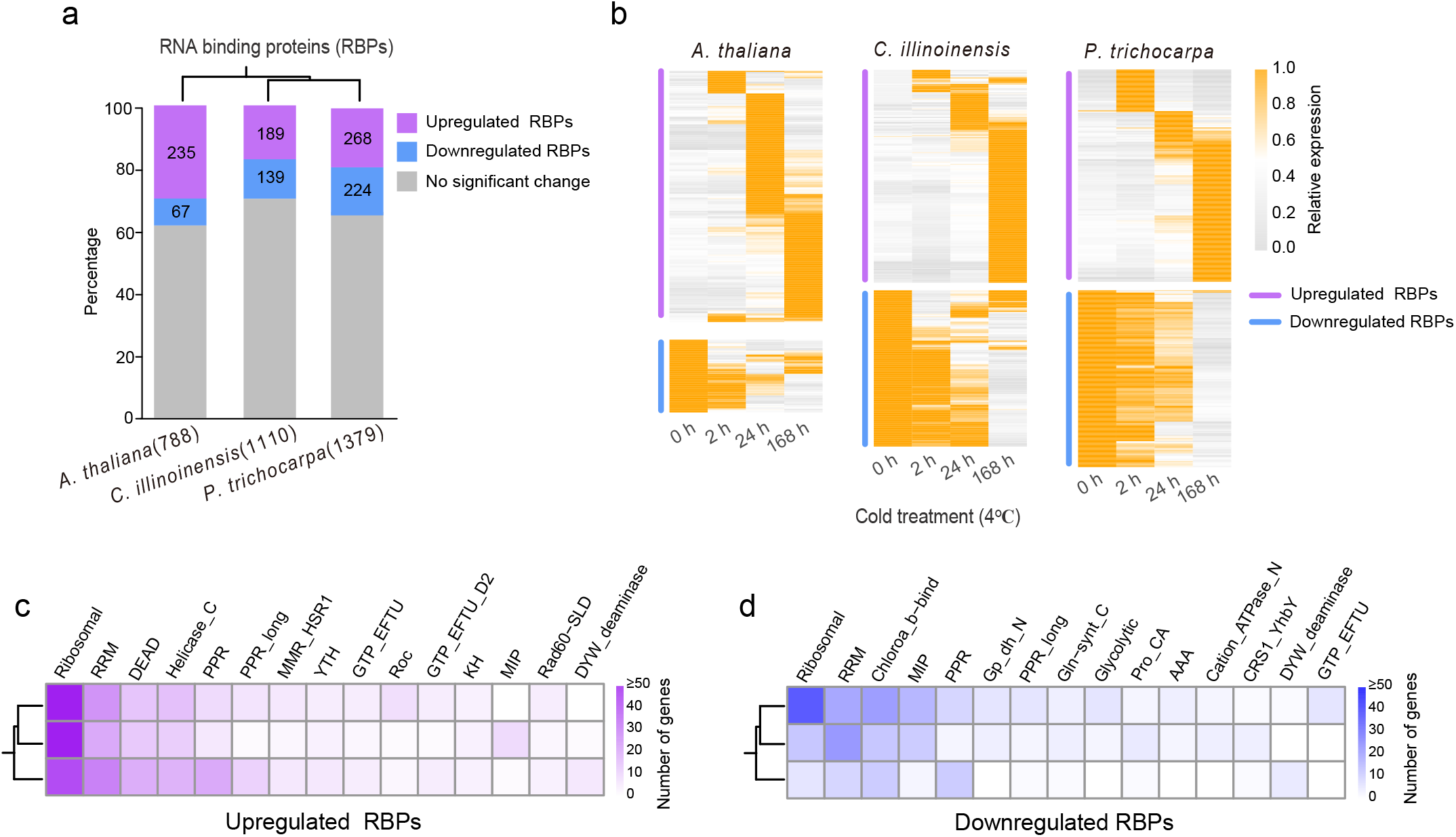
Cold-responsive RNA binding proteins (RBPs) in conserved RBP families. (a) Upregulated and downregulated RBPs under cold stress. Bar percentage indicates the proportion of upregulated, downregulated, and no significant expression changed RBPs to all RBPs. The number in parentheses after the species indicates the number of RBPs in the genome. (b) Expression profiles of up- and downregulated RBPs under cold stress. The expression values of the RBPs at four different timepoints (0, 2, 24, and 168 h) of cold treatments were normalized by row scale. (c-d) The numbers of up- and downregulated RBPs in RBP families. The top 15 up- and downregulated RBP families were selected for visualization.

### Conserved cold-responsive RBP families in rosids

Based on RNA-binding domains, we classified RBP genes into different gene families (**Supplementary Table 4**). Among the top 15 cold-responsive RBP families, ribosomal and RRM are two most prominent families with more genes being upregulated than downregulated under cold stress in all three plant species (**Fig 4c-d**). DEAD and Helicase_C families are mainly enriched in the upregulated RBPs (**Fig 4c**), while Chloroa_b-bind and MIP families are more clustered in the downregulated RBPs (**Fig 4d**). Like TF families, according to the maximum parsimony principle, we suggest that the conserved cold responsivity of the RBP families in three plant species is largely descended from their common ancestor.

### Cold association of transcriptional and post-transcriptional activities in rosids

As described above, thousands of genes were identified as cold-induced DEGs and/or DASGs in three plants (**Fig 1, Fig 3**). Further, we identified that upregulated DEGs show a significant overlapping association with DASGs, that is, 558, 1340, and 1419 up-DE-DASGs respectively in *A. thaliana, C. illinoinensis*, and *P. trichocarpa* (**Fig 5a, Supplementary Table 5**). Similarly, downregulated DEGs also show a significant association with DASGs in the three plant species. The findings suggest that many cold-responsive genes are differentially regulated not only by TFs at transcriptional level but also by RBPs at post-transcriptional level, and these genes might play an important role in rosids under cold stress. In the following sections, we determined whether these genes across the three studied plants are homologous and function in a conserved way.

**Fig 5.**
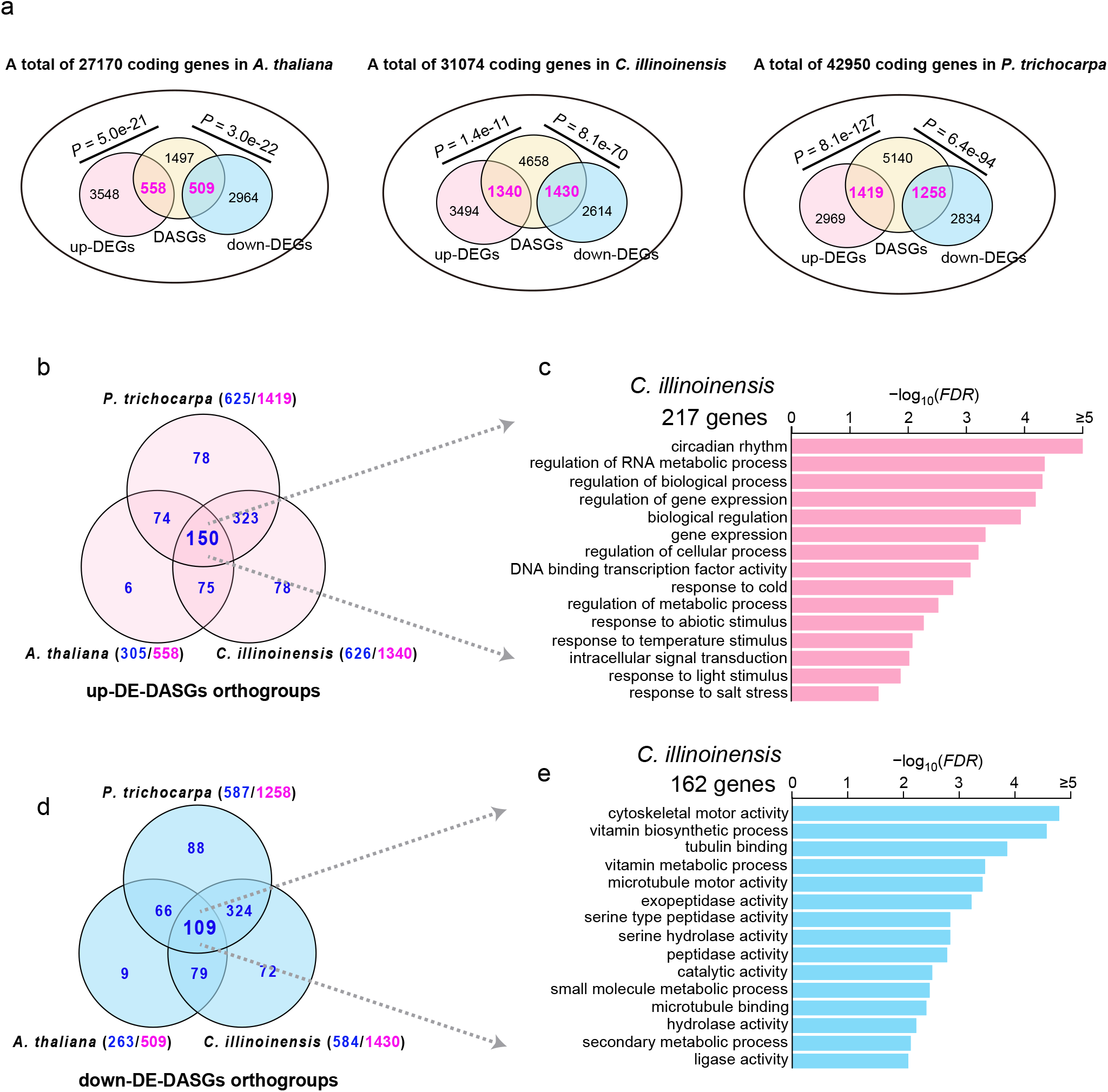
Association of cold-induced DEGs and DASGs and identification of conserved DE-DASG orthogroups in rosids. (a) Overlapping and unique genes between DEGs and DASGs in three species. The overlapping significance was assessed by Fisher’s Exact Test. (b-e) DE-DASG orthogroup identification and enrichment analysis of DE-DASG orthogroup genes in three species. Venn diagram shows overlapping and unique orthogroups of up- (b) and down-DE-DASGs (d) in three species. For each species, the number in purple after the slash indicates the number of up- or down-regulated DE-DASG genes, while the number in blue before the slash indicates the number of the corresponding DE-DASG orthogroups. In the overlapping 150 up- and 109 down-DE-DASG orthogroups, we obtained 217 (c) and 162 (e) *C. illinoinensis* genes representatively for GO term analysis. The same analysis was also performed in *A. thaliana* and *P. trichocarpa* (Supplementary Fig 1). The top 15 significant enriched terms were selected to be shown.

### Identification of 259 conserved orthogroups in rosids

Biased retention and loss of gene duplicates in plants after independent whole-genome and/or small-scale duplications lead to complex and intertwined evolution of co-orthologs and in-paralogs across different species^35,40^. To determine common cold-responsive genes between species, we classified those genes that are differentially regulated at both transcriptional and post-transcriptional levels into orthogroups: sets of homologous genes descended from the recent common ancestor of the plants^41^ (**see Methods and materials**). For example, 558, 1,340, and 1,419 up-DE-DASGs are classified into 305, 626, and 625 orthogroups, respectively in *A. thaliana, C. illinoinensis*, and *P. trichocarpa* (**Supplementary Table 6**). Among these orthogroups, Venn diagram analysis shows a significant overlapping of 150 orthogroups among three plants (**Fig 5b, Supplementary Table 6**). From the 150 orthogroups, we extracted 186 *A. thaliana* genes and performed GO term enrichment analysis (**Fig 5c**). As expected, the genes are mainly enriched in abiotic stresses, such as response to abiotic stresses (salt and cold stress) and circadian rhythm (**Fig 5c**). The GO term enrichments of the counterparts in other two species have similar results (**Supplementary Fig 1**). The same analysis was also conducted for down-DE-DASGs, and we identified 109 overlapping orthogroups with many genes enriched in diverse enzymatic activities (**Fig 5d, 5e, Supplementary Fig 1, Supplementary Table 7**). The findings together show a total of 259 orthogroups (150 + 109) that are regulated at both transcriptional and post-transcriptional levels in a conserved manner in rosids under low temperature, likely suggesting a conserved role of these orthogroups under cold stress.

### Seven striking TF orthogroups involved in circadian rhythms

Among the 259 orthogroups, there are 27 upregulated TF orthogroups (**Supplementary Table 8**) and it is noteworthy that seven of them encode known circadian clock components including *CCA1*/*LHY, RVE4*/*8, RVE7, PRR9, BBX19, COL9*, and *JMJD5* genes (**Table 1**). Our results are consistent with previous reports that *CCA1, LHY* and *PRR9* can be regulated at transcriptional and post-transcriptional levels under cold stress in *Arabidopsis*^31,42^. We showed in our study these circadian genes are also cold-regulated in *C. illinoinensis*, and *P. trichocarpa* and we found that additional circadian clock related genes are differentially cold-regulated at both transcriptional and post-transcriptional levels in rosids (**Table 1**). This explains well aforementioned repeated observations that the biological process of circadian rhythm is enriched (**Fig 1c, Fig 3c, Fig 5c**). The seven orthogroups are upregulated under cold stress in all three plant species, and in contrast, AS types of genes in each orthogroup show a great diversity, which suggest a conservation of gene expression but a rapid evolution of AS events after rosid radiation (**Table 1**). Moreover, MYB is the most prominent TF family including three orthogroups (*CCA1/LHY, RVE7*, and *RVE4/8*). The finding suggests that circadian clock components might play important roles in regulating cold responses in a conserved manner in rosids.

**Table 1.** Cold-responsive circadian-related TFs in rosids.

| Family(orthogroup) | Cold regulation<br>*Ath/Cil/Ptr* | *Ath* (AS-type) | *Cil* (AS-type) | *Ptr* (AS-type) |
| --- | --- | --- | --- | --- |
| MYB (OG0000072) | UP/UP/UP | *CCA1* (A5SS)<br>*LHY*<br>(A3SS,A5SS,RI,SE) | *CIL1205S0016* (A3SS,<br>A5SS, MXE, RI, SE) | *Potri.014G106800* (A3SS,<br>A5SS, MXE, IR, SE) |
| MYB (OG0000110) | UP/UP/UP | *RVE4* (A3SS, SE)<br>*RVE8* (A3SS, IR) | *CIL0982S0042* (A5SS) | *CIL0982S0042* (A5SS) |
| MYB (OG0000043) | UP/UP/UP | *RVE7* (SE) | *CIL1030S0151* (IR)<br>*CIL1245S0064* (IR) | *Potri.004G074300* (IR)<br>*Potri.012G038300* (IR) |
| PRR (OG0000018) | UP/UP/UP | *PRR9* (A5SS) | *CIL0897S0115* (IR)<br>*CIL1083S0010* (IR)<br>*CIL1194S0023* (IR) | *Potri.002G179800* (A3SS,<br>IR, SE)<br>*Potri.014G106000* (A3SS,<br>IR, SE) |
| BBX (OG0000101) | UP/UP/UP | *COL9* (IR) | *CIL1155S0003* (IR) | *Potri.002G214500* (IR)<br>*Potri.014G170600* (IR) |
| BBX (OG0000088) | UP/UP/UP | *BBX19* (IR, SE) | *CIL0978S0047* (IR) | *Potri.001G061800* (A3SS) |
| Cupin (OG0000219) | UP/UP/UP | *JMJD5* (A3SS, IR) | *CIL1465S0014* (IR) | *Potri.001G016200* (IR) |

Note: The short names (*Ath*, *Cil*, and *Ptr*) indicate the species (*A. thaliana*, *C. illinoensis*, and *P. trichocarpa*)

### Cold-responsive genes regulated by circadian clock components

First, we collected experimentally genome-wide regulome maps of *CCA1, LHY, RVE4, RVE7*, and *RVE8* in *A. thaliana* with data from chromatin immunoprecipitation (ChIP-seq)^43^ and DNA affinity purification followed by deep sequencing (DAP-seq)^44^. Subsequently, we developed a stringent pipeline to identify cold-responsive genes regulated by the five TFs (see **Materials and methods**). In brief, we integrated the ChIP-seq and DAP-seq data to identify *A. thaliana* genes bound by either of the five TFs and investigated their orthogroup genes in *C. illinoinensis* and *P. trichocarpa* whose promoters (1-Kb) also have at least one *cis*-regulatory binding site of the TFs. Finally, and of importance, we integrated phylotranscriptomic data to identify those orthologous genes that have DNA binding sites of the five TFs and are differentially cold-regulated in all three plant species. Thus, we designated those orthologous genes as <u>Co</u>nserved <u>COR</u> gene<u>s</u> (CoCORs) targeted by clock components *CCA1, LHY, RVE4, RVE7*, or *RVE8* in rosids (**Fig 6a, Supplementary Table 9**).

**Fig 6.**
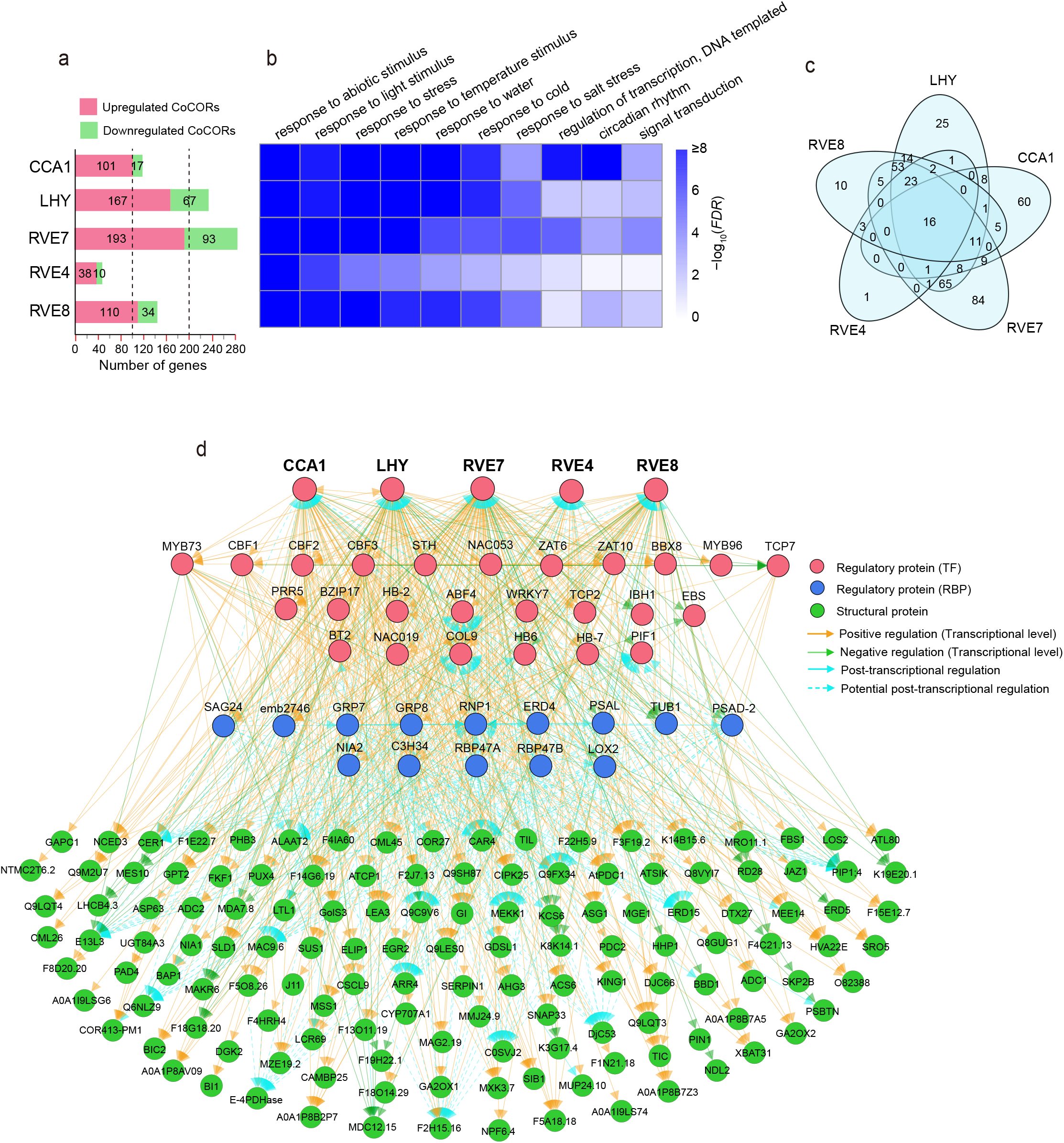
A hierarchical cold-responsive regulatory network launched by circadian clock components in rosids. (a) Numbers of up- and downregulated CoCORs targeted by CCA1, LHY, RVE4, RVE7, and RVE8. These CoCORs were predicted by a stringent pipeline (see Materials and methods). (b) GO enrichment analysis of CoCORs. The top 10 significant GO terms were selected to be shown. (c) Overlapping and unique CoCORs regulated by the five circadian-related TFs. (d) A hierarchical network launched by the five circadian-related TFs. The network is arranged in three layers (TFs, RBPs, and Structural proteins). To improve the robustness of the network, we also integrated regulations of CBFs, ZAT10, HB6, HB-7, NAC053, WRKY7, MYB73, TCP7, and GRP7 in relation to CoCORs. Dotted lines indicate that the two linked genes are co-regulated under cold stress and their potential regulation remains to be determined. Solid arrow lines indicate that additional binding evidence analyzed from ChIP/DAP-seq and iCLIP-seq experiments supports the linked genes.

As shown in Fig 6a, the five circadian components regulate different numbers of CoCORs but with a common pattern of dominant upregulation in three rosids. For instance, as compared to only 17 downregulated CoCORs, there are 101 CoCORs identified to be upregulated by CCA1 (**Fig 6a, Supplementary Table 9**). GO term analysis further shows that these CoCORs are enriched in environmental stresses, such as response to light, cold, and circadian rhythm (**Fig 6b**). Moreover, Venn diagram shows a significance of overlapping CoCORs regulated by the five TFs (**Fig 6c**), and among the overlapping CoCORs, 16 are regulated by all five TFs and 210 are regulated by at least two of the five TFs. The intertwined DNA binding of CoCORs by the five circadian TFs suggests a complex regulatory network launched by circadian clock components.

### A conserved hierarchy of cold-responsive regulatory network in rosids

To construct the network, we employed the CoCORs involved in abiotic stresses that are directly regulated by the five circadian components. Taking account of cold-responsive speeds of the genes under cold treatments at 2, 24, and 168 h and also their regulatory relationships, we constructed a hierarchical regulatory network with three layers launched by the five circadian clock components and elaborated this network as follows.

The first layer is composed of 30 TFs including many well-known cold-regulated genes, such as *CBFs, PRR5, ZAT6, ZAT10, COL9, BT2* etc. In this layer, the five clock components play a dominant role to upregulate the expression of 21 TFs and in turn work together with these upregulated TFs to further regulate the expression of downstream genes (e.g., RBPs and structural genes in the second and third layers). In addition to upregulated TFs launched by circadian clock components, there are four downregulated TFs (*TCP7, EBS, IBH1*, and *PIF1*). This demonstrates a potential negative regulatory loop between differentially regulated TFs for controlling the balance between plant growth and cold tolerance. Evidentially, phytochrome-interacting factor 1 (PIF1) negatively regulates plant freezing tolerance likely through repressing the expression of *CBFs*^45^.

The second layer is composed of 14 RBPs. Among the 14 RBPs, 12 are upregulated by the first layer TFs. These 12 RBPs function in a feedback mechanism by potentially regulating AS of TFs in the first layer and function in regulating AS of the downstream layer genes. For example, by analyzing individual nucleotide resolution crosslinking and immunoprecipitation (iCLIP)-seq experimental data^46^, we found that GRP7 can directly bind to transcripts of *BT2* and *HB6* in the first layer and *COR27, KCS6, BAP1*, COR413-*PM1, ERD15, LTL1, PSBTN*, and *MEE14* in the third layer, and many of these genes undergo differentially AS events under cold stress.

The third layer is composed of 140 *COR*s mainly encoding structural proteins. These genes are differentially regulated in expression and/or alternative splicing by the first and/or second layer regulators. Among the genes, overexpression or knockout of some genes (e.g., *COR47, TIL, SLD1, LOS2*, and *ADC1*/*2*) can affect plant freezing tolerance^47-51^. Altogether, our findings suggest that circadian clock components are actively regulated under cold stress to launch a hierarchy of gene regulatory network to allow adaptation to cold stress in rosids.

## Discussion

When exposed to cold stress, plants undergo a series of molecular changes to protect themselves from chilling and/or freezing damage^3,4^. By high-throughput methods, previous studies provided insights into the reprogramming of gene regulation in response to cold stress^7,8,29,30,32^. However, the studies were generally performed with single plant species and focused on the best-known CBF-dependent pathway genes. Here, we used three plant species (*A. thaliana, C. illinoinensis*, and *P. trichocarpa*) form different rosid lineages which diverged during the early evolution of rosids (∼100 million years ago) and performed RNA-seq experiments with plants subjected to cold stress to investigate conserved regulators and mechanisms underlying rosids in response to cold stress. Over seven thousand DEGs were identified in each species and show common responses of biological processes, such as cold stress process and circadian rhythm (**Fig 1c**). Interestingly, the three plant species show different cold-responsive speeds: an early in *A. thaliana* but a delayed extensive cold-response in *C. illinoinensis* and *P. trichocarpa* (**Fig 1b, 2b**). The difference in cold-responsive speeds maybe due to the different growth characteristics between annual herbaceous plants (e.g., *A. thaliana*) and perennial woody plants (e.g., *C. illinoinensis* and *P. trichocarpa*). Compared to woody plants, herbaceous plants are more sensitive to temperature changes^52^, which might cause an early extensive cold response.

In addition to transcriptional regulation, post-transcriptional regulation is a key process in reprogramming gene expression under cold stress^7,30^. Like DEGs, thousands of DASGs were identified in rosids under cold stress. Interestingly, the number of DASGs in *A. thaliana* is significantly much less than the counterparts in *C. illinoinensis* and *P. trichocarpa* (**Fig 3a**). This difference may be due to shorter introns in *Arabidopsis* than in *C. illinoinensis* and *P. trichocarpa* (**Supplementary Fig 2**), and alternative splicing tends to occur around the exons interrupted by long introns^53,54^. In consistence with DEGs, DASGs show an enrichment of components involved in circadian rhythm (**Fig 3c**). Unexpectedly, enrichments of biological processes related to cold or abiotic stress are not observed for DASGs. This is probably because a limited knowledge of cold-induced DASGs and their genetic roles as compared to an extensive study of cold-induced DEGs. Regarding this, we provide a complete list of cold-induced DASGs in three rosid species (**Supplementary Table 3**) and hope this list will be a potential reference for future research of splicing regulation under cold stress.

Moreover, association analysis of transcriptional and post-transcriptional activities revealed 259 conserved DE-DASG orthogroups in rosids. From DE-DASG orthogroup genes, circadian rhythm is again significantly enriched (**Fig 5c**). Circadian clock system can allocate stress-responsive processes to specific times of the day, which provides a control of over daily energy expenditure and enables plants to grow and reproduce under temperature changes^19,55-57^. Here, we found that some conserved circadian clock components are reprogrammed at both transcriptional and post-transcriptional levels under cold stress in rosids (**Fig 5c**). Among them, MYB is the most prominent TF family including three orthogroups with five *Arabidopsis* genes (*CCA1/LHY, RVE7*, and *RVE4/8*), which likely contribute to the flexibility and controllability of the cold response network. Thus, by integrative omics of phylotranscriptomes and regulome maps (RNA-seq and ChIP/DAP-seq data), we identified their direct targets and constructed a conserved hierarchy of cold-responsive regulatory network launched by the five MYB TFs. The existence of many well-known cold-response genes, such as *CBFs, RBG7/8, ZAT6/10*, and *COR27*, in the network demonstrate a high confidence of the network but also imply an importance of circadian clock for plants in response to cold stress. Altogether, our research shed light on conserved hierarchical cold-responsive network in rosids at transcriptional and post-transcriptional levels via circadian clock components, which provide insights into conserved core regulators and ancestral mechanisms underlying cold-responsive gene regulatory network across rosids.

## Materials and Methods

### Identification and characterization of TFs and RBPs

Proteome sequences and gene annotations of three rosid species (*A. thaliana, C. illinoinensis*, and *P. trichocarpa*) were directly downloaded from the public databases^58,59^. Their divergence times were extracted and assessed from the TimeTree website^34^. TFs and RBPs in the species were obtained through the following strategies. For TFs, we downloaded *A. thaliana* and *P. trichocarpa* TFs from plantTFDB 4.0 and identified TFs in *C. illinoinensis* according to the method as described^60^ (**Supplementary Table 2**). For RBPs, we obtained *A. thaliana* RBPs, which were experimentally confirmed RNA-binding assays^25,39^. By using *A. thaliana* RBPs as queries, we employed BLASTP 2.5.0^61^ to retrieve RBP homologs in *P. trichocarpa* and *C. illinoinensis* (E-value < 1e–10, identity > 60%) (**Supplementary Table 4**).

To identify protein families of TFs and RBPs, we downloaded all Hidden Markov Model (HMM) profiles (>19,000) of protein domains from the Pfam database^62^ and searched them against the proteome sequences of the three plant species by HMMER 3.1b2 with an E-value < 1e–5^63^. Accordingly, we obtained gene families for each TF and RBP (**Supplementary Table 4**).

### Genome-wide analysis of cold-induced DEGs and DASEs

To explore molecular changes of rosids under cold stress, we performed RNA-seq experiments in three above-mentioned plant species with cold treatment (4[) at 0, 2, 24, and 168 hours (h). The experiment and analysis were performed as previously described^64^. Gene expression under normal and cold treatment conditions was represented by Trimmed Means of M value (TMM) calculated from the software StringTie v2.0.3^65^. Subsequently, cold-induced DEGs were defined based on three criteria: at least one sample with TMM greater than 5, more than two-fold change in expression levels before and after cold treatment, and a statistical difference with *P* value < 0.01. Additionally, five types of AS events, including intron retention (IR), alternative 3’ splice site (A3SS), alternative 5’ splice site (A5SS), skipping exon (SE), and mutually exclusive exons (MXE), were identified by rMATS v4.0.2^66^ coupled with CASH v2.2.1^26^. A false discovery rate (FDR) ≤ 0.05 was set as the criterion for identifying cold-induced DASEs between normal and cold treatments.

### GO term enrichments

The GO term annotations of *A. thaliana* genes were directly acquired from The *Arabidopsis* Information Resource (TAIR) database^67^. According to method described by the previous reports^35,68^, the genes in *P. trichocarpa*, and *C. illinoinensis* were annotated with the highest alignment score matches against *A. thaliana* proteins from BLASTP searches (E-value < 1e–5)^61^. The GO term enrichment analysis was performed using the OmicShare tools (www.omicshare.com/tools).

### Identification of DE-DASG orthogroups

To explore whether DE-DASGs across the rosids are homologous, we classified the up- and down-DE-DASGs of three rosids into different orthogroups using Orthofinder v2.3.8^41^ with the parameters (-M dendroblast; -S diamond; -A mafft; -I 1.5). Subsequently, we obtained a total of 1,532 DE-DASG orthogroups and among which, 259 were identified to be present in all three species. In 259 DE-DASG orthogroups, 150 and 109 are respectively upregulated and downregulated under cold stress (**Supplementary Table 6, 7**).

### Identification of CoCORs targeted by circadian clock components

To determine CoCORs regulated by circadian clock components in rosids, we developed the following pipeline. First, we obtained *A. thaliana* ChIP-seq (CCA1) and DAP-seq (LHY, RVE7, RVE4, and RVE8) datasets^43,44^ and performed ChIP/DAP-seq analysis as described^64^. The genes with at least one binding peak of the five TFs in the 1-Kb upstream of translation initiation site (TIS) were considered as direct target genes of the corresponding TFs in *A. thaliana*. Second, we collected *cis*-regulatory binding motifs of CCA1, *LHY, RVE7, RVE4*, and *RVE8*^43,44^ and searched them against the 1-Kb promoter sequences of *P. trichocarpa* and *C. illinoinensis* genes using FIMO (E-value <1e–3)^69^. The genes with at least one *cis*-regulatory binding site for the five TFs were identified as potentially targets of the TFs in *P. trichocarpa* and *C. illinoinensis*. Subsequently, all coding genes in the three species were classified into orthogroups according to their homology. The orthogroup genes that have binding sites of the five TFs and are differentially regulated after cold stress in all three rosids are defined as CoCORs. Finally, a total of 101/17, 167/67, 193/93, 38/10, and 110/34 CoCORs (**Fig 6a, Supplementary Table 9**) were identified as to be up/down-regulated by CCA1, LHY, RVE7, RVE4 and RVE8, respectively.

### Statistical analysis

The significance of enrichment analysis between gene groups was calculated by Fisher Exact test through R programming.

## Acknowledgments

We thank the members from Dr. Wenwu Wu’s laboratory in Zhejiang A&F University for suggestions to improve the quality of the study. This work was supported by the National Natural Science Foundation of China (grant number 31871233).

## Contributions

W.W., Z.J., and J.X. designed the research; L.G. and X.Z. performed the study; S.W. did the RNA-seq experiments of the plants and Y.X. performed the RNA-seq analysis; L.G., W.W. and Z.J. wrote and edited the manuscript.

## Data availability statement

The RNA-seq datasets of the three plant species including *A. thaliana, C. illinoinensis*, and *P. trichocarpa* under cold stress have been deposited to NCBI BioProject (Accession: PRJNA767196) and CNGBdb (Accession: CNP0002243).

## Supplementary information

Supplementary Fig 1. The GO term enrichments of conserved up- and down-DE-DASGs in *P. trichocarpa* and *C. illinoinensis*.

Supplementary Fig 2. Alternative splicing seems to occur in the genes with longer introns.

Supplementary Table 1. RNA-seq values of DEGs in rosids at four timepoints (0, 2, 24, and 168 h) of cold treatments (4[).

Supplementary Table 2. TF family genes and expression in rosids under cold stress. Supplementary Table 3 A list of cold-induced DASGs and their AS types in rosids. Supplementary Table 4. RBP family genes and expression in rosids under cold stress. Supplementary Table 5. The detailed information of DE-DASGs.

Supplementary Table 6. The orthogroups of up-DE-DASGs in rosids. Supplementary Table 7. The orthogroups of down-DE-DASGs in rosids. Supplementary Table 8. Details of 27 up-DE-DASG TF orthogroups in rosids.

Supplementary Table 9. The predicted CoCORs targeted by clock components of CCA1, LHY, RVE4, RVE8, and RVE7 in rosids.

