## Supplementary Figure 1 for "Integrative multispecies omics reveals a hierarchy of cold-responsive regulatory network launched by circadian components in rosids"

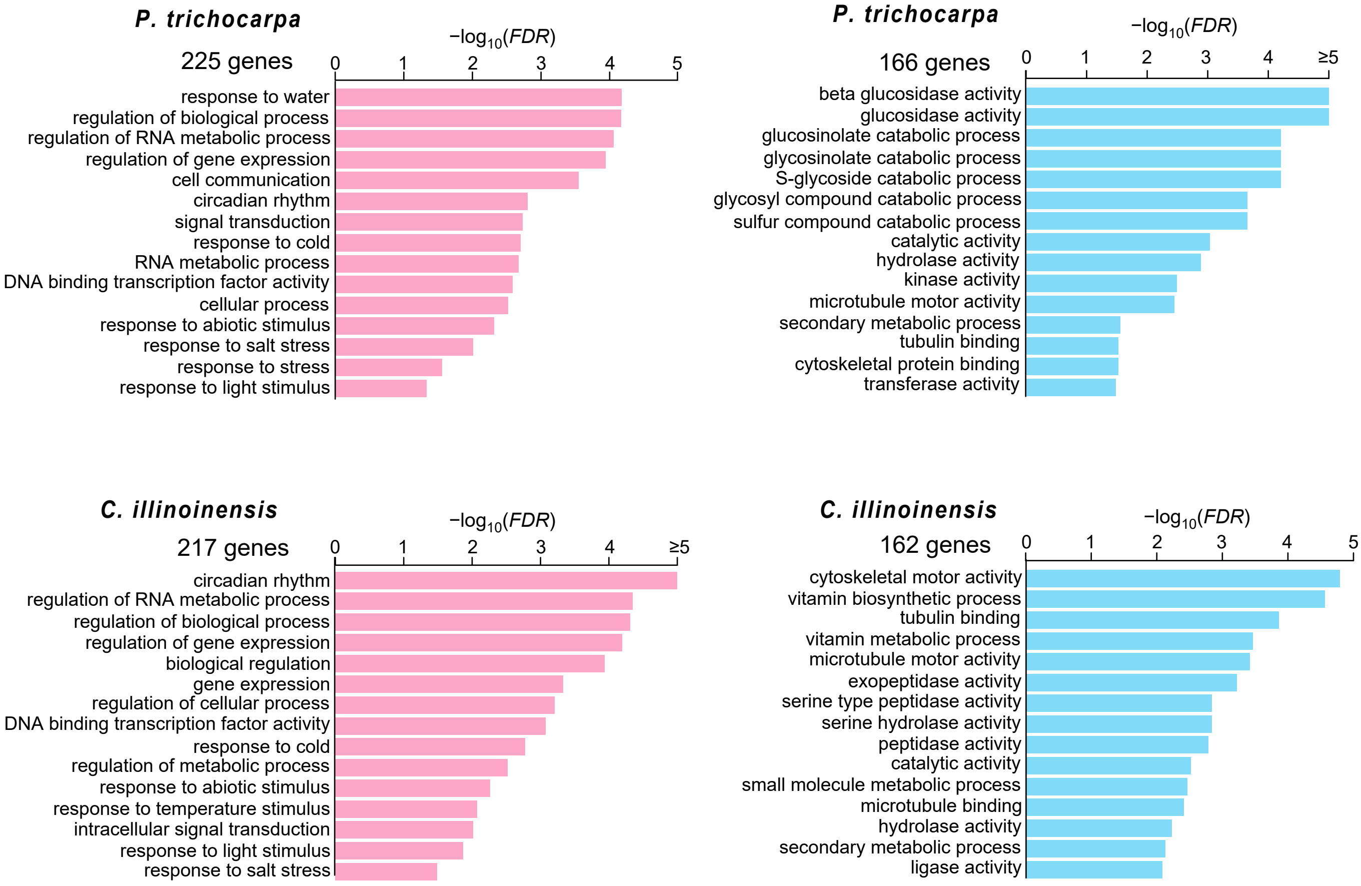


**Supplementary Fig.1. The GO term enrichments of conserved up-DE-DASGs (red) and down-DE-DASGs (blue) in *P. trichocarpa* and *C. illinoinensis.***
