## Supplementary Figure 2 for "Integrative multispecies omics reveals a hierarchy of cold-responsive regulatory network launched by circadian components in rosids"

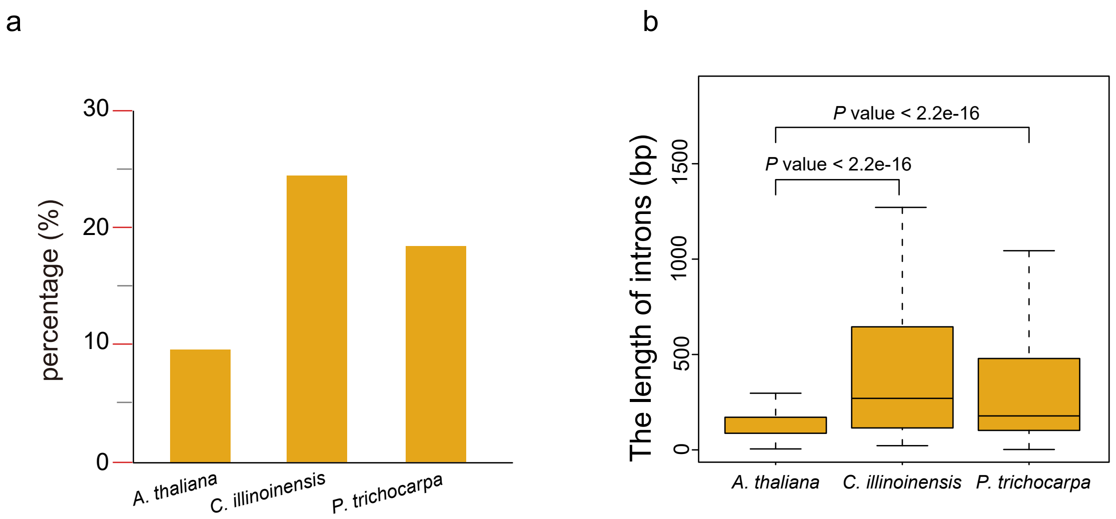


**Supplementary Fig.2. Alternative splicing seems to occur in the genes with longer introns.** (a)The percentages of DASGs to all genes in the plants. (b) The length of all introns in three rosids. The difference was assessed by the nonparametric Mann-Whitney test.
